# Shared associations identify causal relationships between gene expression and immune cell phenotypes

**DOI:** 10.1101/2020.09.19.304832

**Authors:** Christiane Gasperi, Sung Chun, Shamil R. Sunyaev, Chris Cotsapas

**Affiliations:** Department of Neurology, Yale School of Medicine, New Haven CT USA; Division of Genetics, Brigham and Women’s Hospital, Boston, MA USA; Department of Biomedical Informatics, Harvard Medical School, Boston, MA, USA; Department of Genetics, Yale School of Medicine, New Haven CT USA

## Abstract

Genetic mapping studies have identified thousands of associations between common variants and hundreds of human traits. Translating these associations into mechanisms is complicated by two factors: they fall into gene regulatory regions; and they are rarely mapped to one causal variant. One way around these limitations is to find groups of traits that share associations, using this genetic link to infer a biological connection. Here, we assess how many trait associations in the same locus are due to the same genetic variant, and thus shared; and if these shared associations are due to causal relationships between traits. We find that only a subset of traits share associations, with most due to causal relationships rather than pleiotropy. We therefore suggest that simply observing overlapping associations at a genetic locus is insufficient to infer causality; direct evidence of shared associations is required to support mechanistic hypotheses in genetic studies of complex traits.

## Introduction

Genetic mapping studies have identified thousands of associations between common variants and hundreds of human traits. Uncovering the mechanisms that underlie these traits requires understanding the molecular, cellular and physiological events altered by causal genetic variants. Incomplete fine mapping due to linkage disequilibrium and the possible action of causal variants across diverse cell types, contexts and genes currently limits our ability to infer the mode of action of causal variants, and hence the biology underlying traits. Experimentally testing multiple such mechanistic hypotheses across thousands of associations rapidly becomes a problem of scale; we thus need principled approaches to generating and testing such mechanistic hypotheses.

We and others have suggested such an approach, building on the concept of pleiotropy. The molecular and cellular events altered by causal genetic variants are, by definition, also genetic traits, and they must be associated with the same variant. To link traits together and thus form mechanistic hypotheses, one can thus look for shared genetic associations between traits^1–4^. Such sharing is often defined as two traits associated with variation in the same general genome region, often within an arbitrary window of physical distance. A more robust alternative is to identify pairs of traits that share an underlying causal effect, rather than a shared genomic segment, and several methods have been developed to this end^5–12^.

We have reported a relative paucity of overlaps between expression quantitative trait loci (eQTLs) and disease risk associations^12^. This result appears paradoxical given the strong enrichment of risk heritability in gene regulatory regions^13–15^, which suggests that the majority of risk effects should alter gene regulation, and therefore expression. This paucity may be because we are not interrogating the right cell types, or the right physiological or stimulation conditions for those cells; or it may be that we lack power to detect such overlaps, though our simulations suggest the latter is not a major factor^3,12^.

It is tempting, therefore, to assume that requiring a demonstration of shared association between traits is overly stringent, and simply identifying associations to the same region is a more productive approach. A further temptation is then to assume causality, especially when the two traits are drawn from different levels of physiology: it is natural to assume that a gene expression trait is causal for disease risk rather than the other way around, for example. This assumption of causality is made implicitly when pairs of associations are used to propose mechanistic hypotheses of pathophysiology. However, it is also possible that the two traits are either associated to different variants in the same locus, or to a single pleiotropic variant and otherwise share no biological underpinning (horizontal pleiotropy). To our knowledge, there is no *a priori* way to set a prior expectation for causality or pleiotropy. How useful, therefore, is it to identify shared effects between two traits, rather than simply identify associations to the same broad locus?

Here, we answer this question by first assessing how many associations to different traits in the same locus are due to the same underlying effect, and thus shared; and if these associations shared between traits are likely due to a causal relationship between these traits, or if horizontal pleiotropy is widespread. We compare 164 distinct immune cell phenotypes from the Milieu Intérieur project^16^ to gene expression traits in monocytes, neutrophils and T cells from the BLUEPRINT consortium^17^. We first select pairs of immune traits and gene expression traits with associations at the same genetic locus, and then identify which of these pairs share an association and which are associated with different variants in close proximity. We find that trait pairs with shared genetic associations are more likely to share a broader genetic correlation and are more likely to share a causal relationship, as assessed by two Mendelian randomization approaches. Our results show that a substantial proportion of shared associations between traits is likely to be due to causal relationships. We therefore suggest that simply observing associations of different traits to the same genetic locus is insufficient to infer causality, and direct evidence of shared association must be the minimum evidence required to link traits.

## Methods

A schematic overview of our analysis is presented in Figure S1. Unless otherwise specified, all analyses were carried out with R v3.4.1^18^.

### Milieu Intérieur project immune phenotype data processing

We obtained imputed genotype data and flow cytometry measurements for 166 immune phenotypes (75 innate immune cell parameters, 91 adaptive immune cell parameters; see Table S1) for 816 healthy, unrelated people of Western European ancestry from the Milieu Intérieur project^16^. We removed two phenotypes due to the low number of non-zero values (Figure S2a and S2b). We found that all but one of the remaining phenotypes were not normally distributed (Shapiro-Wilk test), so we performed rank-inverse transformation on all phenotypes. After this transformation, four phenotypes still showed evidence of non-normal distribution. From visual inspection, two of these phenotypes - the number of founder B cells and the number of HLA-DR^+^/CD4^+^ EMRA T cells (Figure S2c and S2d) - appeared not to be detectable in a substantial subset of individuals. We therefore reduced these to binary detected/not detected phenotypes. For each phenotype, we defined the detection threshold as the point of the quantile-quantile plot with the highest slope (Figure S3). The other two phenotypes had approximately normal distributions despite the Shapiro-Wilk test (Figure S2e and S2f), so we did not modify them further.

All 816 individuals had been genotyped using the HumanOmniExpress-24 BeadChip and quality control (QC) and genotype imputation has been performed as described in the original publication^16^, yielding a final data set of 5,699,237 SNPs with an IMPUTE Score > 0.8 and a minor allele frequency (MAF) > 0.05. We performed additional QC, removing all individuals with excess heterozygosity of more than five standard deviations from the sample mean (n=6), one sample from each pair showing cryptic relatedness (identical by descent (IBD) > 0.1875, n=3) and population outliers with a distance in the first four principal components of more than 4 standard deviations (n=10). These QC steps were all based on a set of variants with MAF > 0.05, genotyping rate > 98% and a Hardy-Weinberg equilibrium (HWE) test *p*-value > 1×10^-3^ and pairwise linkage disequilibrium (LD) < 0.2. From the complete dataset, we then removed all variants out of Hardy-Weinberg equilibrium (p < 1×10^−5^) and MAF < 0.05, and insertions, deletions and multiallelic variants. Our final dataset was thus 5,231,477 variants across 797 individuals.

### BLUEPRINT expression QTL data processing

We obtained RNAseq data for naive CD4^+^ T cells (169 individuals), CD14^+^ monocytes (193 individuals) and CD16^+^ neutrophils (196 individuals) from the BLUEPRINT consortium, ascertained to be free of disease and representative of the United Kingdom (UK) population^17^. We downloaded FASTQ files and used the GTEx pipeline for RNA-seq alignment, quantification and quality control (https://www.gtexportal.org/, Analysis Methods for V8). Briefly, we performed alignment to the human reference Genome CRCh38/hg38 using STAR v2.5.3a^19^, based on the GENCODE 26 annotation and gene-level quantification with RNA-SeQC v1.1.9^20^. We produced read counts and “transcript per million” (TPM) values as described in the GTEx pipeline. We then selected genes with expression values of > 0.1 TPM and ≥ 6 reads in at least 20% of the samples and normalized between samples using “trimmed mean of M-values” (TMM) as implemented in *edgeR*^21^. We then normalized expression values across samples using an inverse normal transformation. All samples had at least 10 million unique reads. From the BLUEPRINT data release, we obtained genotype data for all individuals for 7,008,524 variants acquired by whole genome sequencing. Sequencing, alignment, variant calling and quality control had been performed as described in the original publication^17^. We additionally filtered out insertion/deletion and multiallelic variants, and all variants with a MAF < 0.05 and a Hardy-Weinberg equilibrium chi-square *p*-value of < 1×10^−5^. We performed sample QC as described above for the MIP dataset, which did not lead to the removal of any individuals, yielding a final genotype data set of 197 individuals and 4,853,096 single nucleotide polymorphisms (SNPs) (GRCh37 build). In total, we found 4,355,418 SNPs present in both the BLUEPRINT and the Milieu intérieur project data sets.

Both the BLUEPRINT and the Milieu intéreur project genotype data sets were available in the GRCh37 build, but version 8 of the GTEx pipeline for RNAseq alignment and quantification uses GRCh38. We reconciled the different genome builds by back-lifting the RNAseq data to GRCh37, determining the transcription start site for each gene with R/BiomaRt v2.34.3^22,23^.

### Association analyses

We performed all association regression analyses with plink v1.9^24^, assuming an additive model of inheritance for all variants. We adjusted all regression analyses on the immune phenotypes from the MIP data set for age, sex, as well as two environmental factors - smoking (0=Non-smoker, 1=Ex-Smoker, 2=Smoker) and latent CMV infections (CMV serology 0=negative, 1=positive) - as these have been identified as the main non-genetic factors affecting immune phenotype variation in the original study^16^. Additionally, we corrected the regression models for the top five principal components to adjust for population stratification. For the association analyses on gene expression data (eQTL analyses) we included age, sex, the first five principal components as well as 30 PEER factors^25^ (calculated as described in the GTEx pipeline) as covariates. We used the same covariates to generate permutation data for JLIM.

### Identifying immune and gene expression trait associations in the same locus

JLIM compares association data for a primary trait to association data for a secondary trait. In all analyses, we use the Milieu Intérieur immune phenotypes as primary traits and BLUEPRINT gene expression traits as secondary. We thus first identify potential associations in the immune phenotypes and then look for overlapping BLUEPRINT eQTLs.

We first identified all independent immune phenotype associations by selecting lead SNPs that i) had suggestive levels of association (p < 1 ×10^−5^); ii) are not within 100 kilobases from another lead SNP; and iii) are not within 500 kilobases from another lead SNP *and* in LD (r^2^ > 0.2) with another lead SNP. To identify conditionally independent associations, we performed stepwise conditional association analyses for all markers within 200 kilobases of each lead SNP. At each step, we identified the most associated SNP not in LD with any other lead SNP (r^2^ < 0.2); if this SNP had p < 1 ×10^−3^, we added it to the model and repeated the analysis until no independent SNP satisfied the *p*-value threshold. For each conditionally independent signal, we then calculated residual association statistics, where we condition on all other independent effects in a locus. All association signals with a lead SNP with an association p < 1 ×10^−5^ were carried forward to subsequent analyses. These represent strong independent associations, with any residual weak effects removed (identified by the more lenient p < 1 ×10^−3^ threshold).

We next identified *cis*-eQTLs overlapping immune phenotype associations. We adopted the GTEx definition of a *cis*-eQTL being within 1 megabase of the transcription start site of the gene. We looked for conditionally independent immune phenotype associations within 200 kilobases of each lead SNP above; we therefore identified all genes with a transcription start site (TSS) within 1 megabase of each lead SNP (R/BiomaRt v2.34.3, Ensembl build 37). We then looked for cis-eQTL associations for each such gene in T cells, monocytes and neutrophils, independently. For each identified eQTL (p < 1 ×10^−3^), we then performed stepwise conditional association analyses as described above, for all SNPs within 1.2 megabases of the TSS (Figure S4). We chose this distance so any effects overlapping the lead SNP window are conditionally independent.

Due to the smaller sizes of the gene expression traits we limited iterations to a maximum of three independent signals per locus. As above, we then calculated residual association statistics for each independent eQTL effect in each of the three cell types. For each of the genetic loci associated with immune phenotypes we then selected all gene expression association statistics with a lead SNP with an association p < 1 ×10^−3^ at the respective genetic locus.

### Identifying shared associations between immune and gene expression traits

We tested for shared effects between immune and gene expression trait pairs with JLIM v2^12^. Given genotype-phenotype associations for two phenotypes in different cohorts in the same locus, JLIM assesses the likelihood of the joint model that variant *i* is causal in one trait and variant *j* in another trait, over some number of variants observed in two distinct cohorts. If this joint likelihood is maximal when *i* ≠ *j* we can infer the presence of a single, shared effect driving both associations. Conversely, when the likelihood is maximal when *i* ≠ *j* we can infer that the observed associations are due to different underlying effects. JLIM assumes that only one causal variant for each of the tested traits is present in the analyzed window.

For each trait pair, we used the immune phenotype as the primary trait and the gene expression trait as secondary. We used the 404 non-Finnish European samples from the 1000 Genomes Project (phase 3, release 2013/05/02) as an external LD reference panel. We permuted the secondary trait for each pairwise comparison 100,000 times to obtain empirical significance levels, and used a false discovery rate (FDR) < 0.05 as a significance threshold.

We compared 16,652 unique combinations of immune phenotype and gene expression trait in one of three cell types across 1,199 genetic loci. These pairs encompassed all 164 immune traits and 7,060 genes with eQTLs in at least one of the three BLUEPRINT cell types. As we considered up to three conditionally independent associations per gene and locus, we made a total of 22,379 comparisons.

### Calculating polygenic risk scores within trait pairs

We used polygenic risk scores (PRS) to assess the global genetic overlap between immune/gene expression trait pairs, beyond the shared associations identified by JLIM. For each pair, we selected independent variants associated with the expression trait at some threshold and then calculated PRS for all individuals in the immune trait cohort, using PRSice v2.2.11.b^26^ with default parameters for clumping and an additive genetic model. A PRS *Ŝ_i_* for the *i*th individual over *m* independent SNPs is defined as:

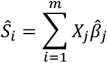

where *X_j_* is the number of minor alleles carried at the *j*th SNP, and 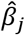 is the eQTL effect size for the *j*th SNP^27^. To account for shared effects between some trait pairs but not others, we condition eQTL traits on the variant with the strongest JLIM *p*-value (even if not significant), and use the now conditionally independent eQTL data for the PRS calculation. We used ten different significance thresholds to select these SNPs: 1 ×10^−2^, 1 ×10^−3^, 1 ×10^−4^, 5×10^−5^, 1 ×10^−5^, 5×10^−6^, 1 ×10^−6^, 5×10^−7^, 1 ×10^−7^, and 5×10^−8^. We then calculated the proportion of immune phenotype variance *(F^2^)* explained by these PRS and their empirical significance, also using PRSice.

We then compared PRS results between trait pairs with a shared effect and trait pairs with no sharing, using two approaches. We compared the proportion of immune phenotype variance explained (i.e. the *F^2^* values) with the Mann-Whitney-Wilcoxon test, and the correlation between JLIM *p-*values and PRS *F^2^* with univariate linear regression.

### Mendelian randomization analyses

We used two Mendelian randomization (MR) approaches to assess evidence that gene expression traits are causal for the immune phenotype traits for which they share an association. First, we used two-sample Mendelian randomization (TSMR)^28,29^, as implemented in the *TwoSampleMR v0.4.25* R package, using inverse variance weighting of effect sizes. As instruments, we selected all independent SNPs associated with the gene expression trait with an association *p*-value < 1 ×10^−5^.

We also used transcriptome-wide summary statistics-based Mendelian Randomization (TWMR), an extension of TSMR^30^. As described by Porcu *et al,* we first identified all variants associated with the gene expression trait in each trait pair in each cell type (conditional association *p* < 1×10^−3^). We then identified all genes within 1 megabase of the gene’s TSS, and selected all SNPs associated with the expression levels of any of these genes. We then removed genes with highly correlated expression values to the original gene (*r^2^* > 0.2), and selected pairwise-independent SNPs from the remaining list (pairwise LD *r*^2^ < 0.1). We used the resulting set of variants as instruments in a multivariate MR model to estimate the causal effect on the immune phenotype.

We compared causality estimates from both methods between trait pairs with a shared effect and trait pairs with no sharing. We compared estimated causal effect sizes with the Mann-Whitney-Wilcoxon test; and, as a continuous measure, the correlation between JLIM *p*-values (strength of evidence of shared effect) with the estimated causal effect sizes and corresponding *p*-values using univariate linear regression.

### Data availability

No data were generated beyond the publicly available datasets used.

### Code availability

Details about software and algorithms used in this study are given in the “Methods” section. No customized code or algorithm deemed central to the conclusion was used.

## Results

### Immune phenotype loci harbor eQTLs but do not share associations with them

To identify effects shared between immune and gene expression phenotypes, we first identified associations for each of the 164 immune phenotypes included in the Milieu Intérieur Project data. JLIM, our joint likelihood mapping method, detects shared associations based on patterns of LD, so we excluded the major histocompatibility complex (MHC) locus on chromosome 6, where LD structure is particularly complex. We found 1,379 distinct non-MHC loci (200 kilobases windows centered on the most associated variant) with evidence of independent association to at least one immune phenotype (*p* < 1×10^−5^; 32 loci at *p* < 5×10^−8^). In 83/1,379 loci, we found more than one independent effect for the same immune phenotype (conditional *p* < 1 ×10^−5^, 6% of loci).

We next looked for genes whose expression could plausibly be influenced by the same genetic effect in these loci. We found 14,634 genes with a transcription start site within 1 megabase of the lead variants, with at least one gene in 1,318/1,379 (95.6%) of the loci. We found that 7,365/14,634 (50.3%) genes in 1,201/1,318 (91.1%) of loci have an eQTL in at least one immune cell type profiled by the BLUEPRINT Consortium *(p* < 1×10^−3^). Most of these 7,365 genes were influenced by more than one conditionally independent eQTL within a 1.2 megabase window around the TSS: 3,762/5,748 (65.4%), 3,476/5,004 (69.5%), and 4,146/5,906 (70.2%) in T cells, neutrophils and monocytes, respectively (Figure S5). We included all these effects in our subsequent analyses, to make sure we capture all possible gene expression effects. Thus, consistent with previous reports across a spectrum of human traits^12,31^, most loci associated with an immune phenotype also harbor at least one *cis*-eQTL, and many eQTLs have conditionally independent effects^32,33^.

We used JLIM to assess if the immune phenotypes and nearby eQTLs were driven by the same underlying genetic effect, indicating shared mechanisms. After filtering, we compared 22,379 pairs of conditionally independent associations representing all 164 immune traits and 7,060/7,365 (95.6%) of genes, at 1,199/1,379 (86.9%) of the discovered loci. We found evidence for a shared underlying causal variant for 207/22,379 (0.9%) pairs, involving 92/164 (56.1%) immune phenotypes and 127/7,060 (1.8%) distinct genes, at an FDR < 0.05. Thus, though we test the vast majority of cases where immune phenotypes and eQTLs overlap, we find limited statistical evidence for shared effects between them. This is consistent with our previous observations of limited sharing between autoimmune disease associations and *cis-* eQTLs^12^. Also similar to our previous findings, we observe that the power to detect shared association thus depends in part on statistical power in the secondary trait cohorts (Figure S6).

Some of our results highlight clear relationships between traits: for example, we see a shared effect between expression level of *SELL* and the level of its protein product L-selectin (CD62L; Figure 1). We find the eQTL in all three BLUEPRINT cell types, and the immunological trait association in both neutrophils and eosinophils. As expected, the allele associated with increased *SELL* transcript abundance in neutrophils and monocytes is also associated with CD62L intensity in both neutrophils and eosinophils (Figure 1 and Figure S7). However, the same effect decreases *SELL* transcript abundance in T cells. We find strong evidence that the T cell eQTL and CD62L intensity trait association are shared, suggesting a distinct mode of regulation in different cell types (for example, different factors binding to the same regulatory element in different cell types, with both types of interactions influenced by the same genetic variant).

**Figure 1:**
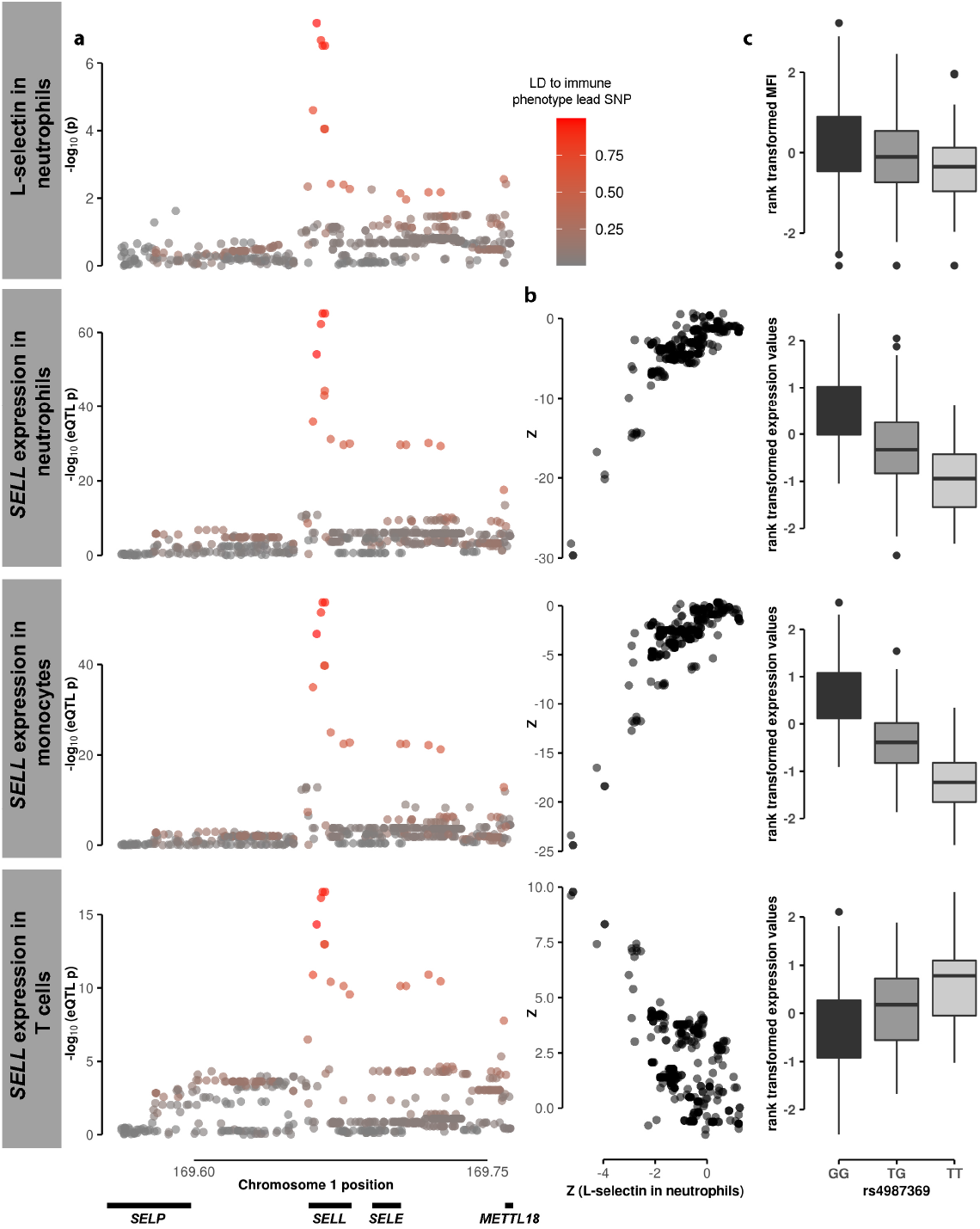
Shared genetic association on chromosome 1 for L-selectin (CD62L) in neutrophils and SELL expression in neutrophils, monocytes and T cells. The association signal of the mean fluorescence intensity (MFI) of L-selectin (CD62L) in neutrophils at a genetic locus on chromosome 1 is consistent with the association signal to *SELL* expression in neutrophils, monocytes and T cells at the same genetic locus (a). The association Z statistics of the mean fluorescence intensity (MFI) of L-selectin in neutrophils and *SELL* expression in the three different cell types are strongly correlated (b). (c) shows the MFI of L-selectin and *SELL* expression in the three different cell types (both rank transformed) per genotype of the lead variant rs4987369.

We found that many shared effect *cis*-eQTLs are not in the same immune cell subpopulation as their cognate immune phenotype. In some cases we did not have expression data for the subpopulation in which an immunophenotype was measured. We found, for instance, a shared effect between the expression level of *CR2* in T cells and the level of its protein product, complement receptor type 2 or CD21 in multiple B cell populations (Figure S8). CD21 is the route through which Epstein-Barr virus infects B cells, and there is more recent evidence that this is also the mechanism of T cell infection^34^. This may therefore be a constitutive *CR2* eQTL, and may have a bearing on susceptibility to EBV infection.

We also found more complex patterns of sharing between cell types. We found several examples where a T cell immune parameter had a shared effect with an eQTL in monocytes or neutrophils, but the gene was either not expressed in T cells or there was no evidence of a shared eQTL in our data (Figure 2). This suggests the possibility that changes to gene expression in one cell type have effects on population number and behavior of another cell type, consistent with the complex and dynamic interplay between immune cell subpopulations.

**Figure 2:**
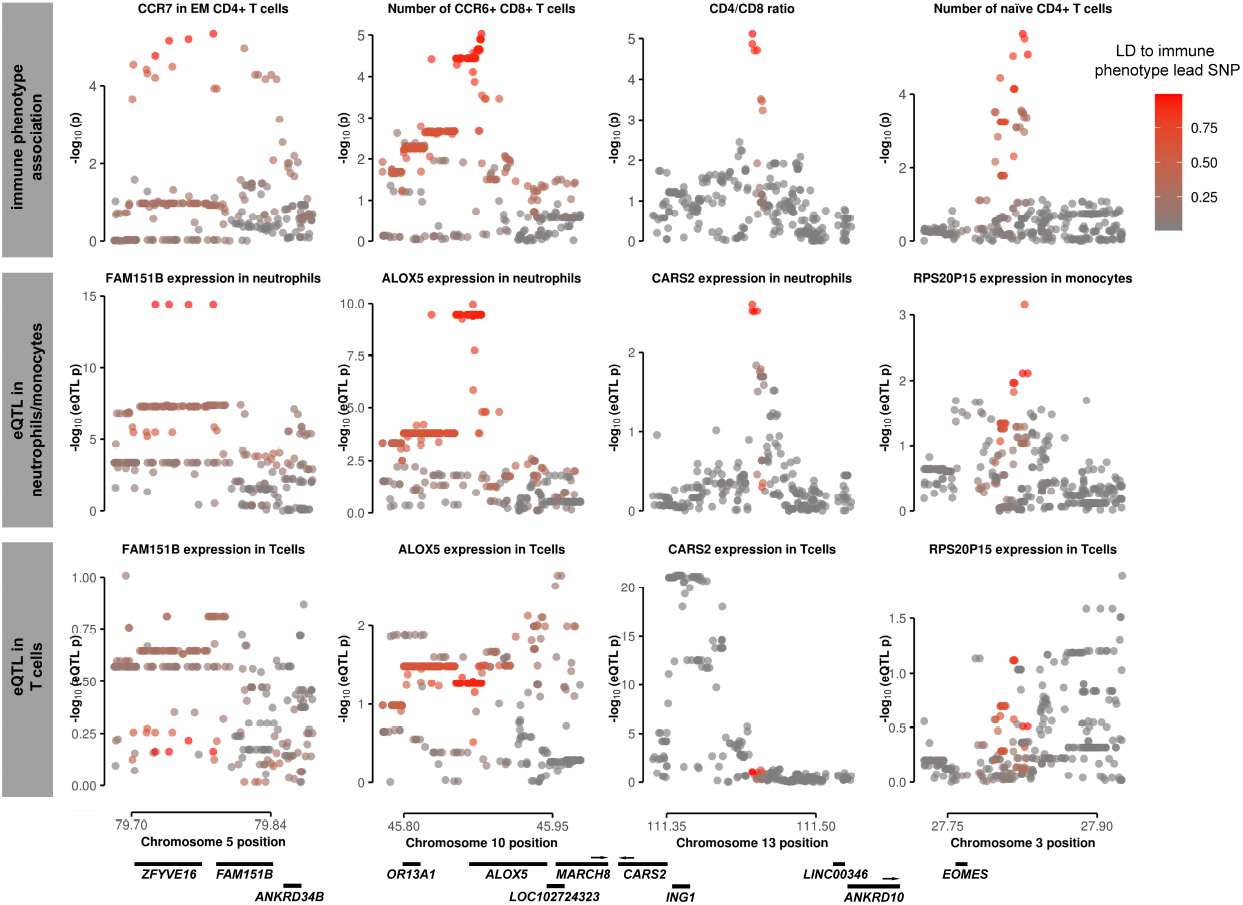
Shared genetic effects between gene expression traits in neutrophils or monocytes, but not in T cells, with immune phenotypes measured in T cells. Shared association signals for different T cell immune phenotypes (first row) and gene expression traits in neutrophils or monocytes (second row). There were no consistent association signals for these gene expression traits in T cells (third row).

### Trait pairs sharing associations show broader genetic correlation

Our broader goal is to establish whether shared associations can identify traits that are causally linked. We therefore sought to distinguish between horizontal pleiotropy, where the same variant influences two otherwise unrelated phenotypes; and mediation, where one phenotype is causal for the other. In horizontal pleiotropy, there should be no further genetic relationship between the traits. Conversely, in the case of mediation we expect the two traits to share genetic architecture more broadly, as perturbation of the intermediate trait should have an effect on the outcome trait^35–37^. Therefore, to establish if traits with shared associations are more likely to be causally related, we first assess evidence for shared heritability between them, and then directly assess evidence for mediation.

To assess evidence for shared heritability between immune-expression phenotype pairs with a shared association, we asked if PRS for the gene expression trait in each pair predicts the immune trait. We compared 190 trait pairs with evidence of shared effects (92 immune and 127 expression traits in 116 loci) to 16,462 trait pairs that do not share the same causal variant per our JLIM analysis. For each trait pair, we calculated a genome-wide PRS for the gene expression trait, and determined the variance of the paired immune phenotype explained by that PRS (as 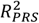). We reasoned that if traits with a shared association are more likely to share heritability more broadly, we should see more variance explained in the 190 trait pairs than in the 16,462 that do not pass our JLIM analysis. We knew, however, that the presence of a shared association would bias this analysis in favor of our expected outcome, because we would be including a known positive association to the immune traits only in the 190 pairs. We accounted for this bias by performing conditionally independent association testing for the main *cis*-eQTL effect.

We found that the variance of an immune trait explained by gene expression PRS was higher when the two traits shared an association than when they did not share one, even though we removed the effect of the shared association (Mann-Whitney-Wilcoxon p < 0.05; Table 1 and Figure 3). We found that this was generally true across a range of thresholds for selecting variants to include in the PRS calculation^27^. We also found that in cases where an immune trait shared an association with more than one eQTL, including PRS for all shared eQTLs explained more variance than any single expression trait alone (Figure S9). Our analysis is conservative, as we condition on the variant with the strongest evidence of shared association; as expected, including lead cis-eQTL effects in the PRS calculations shows an even more extreme difference (Table S2).

**Figure 3:**
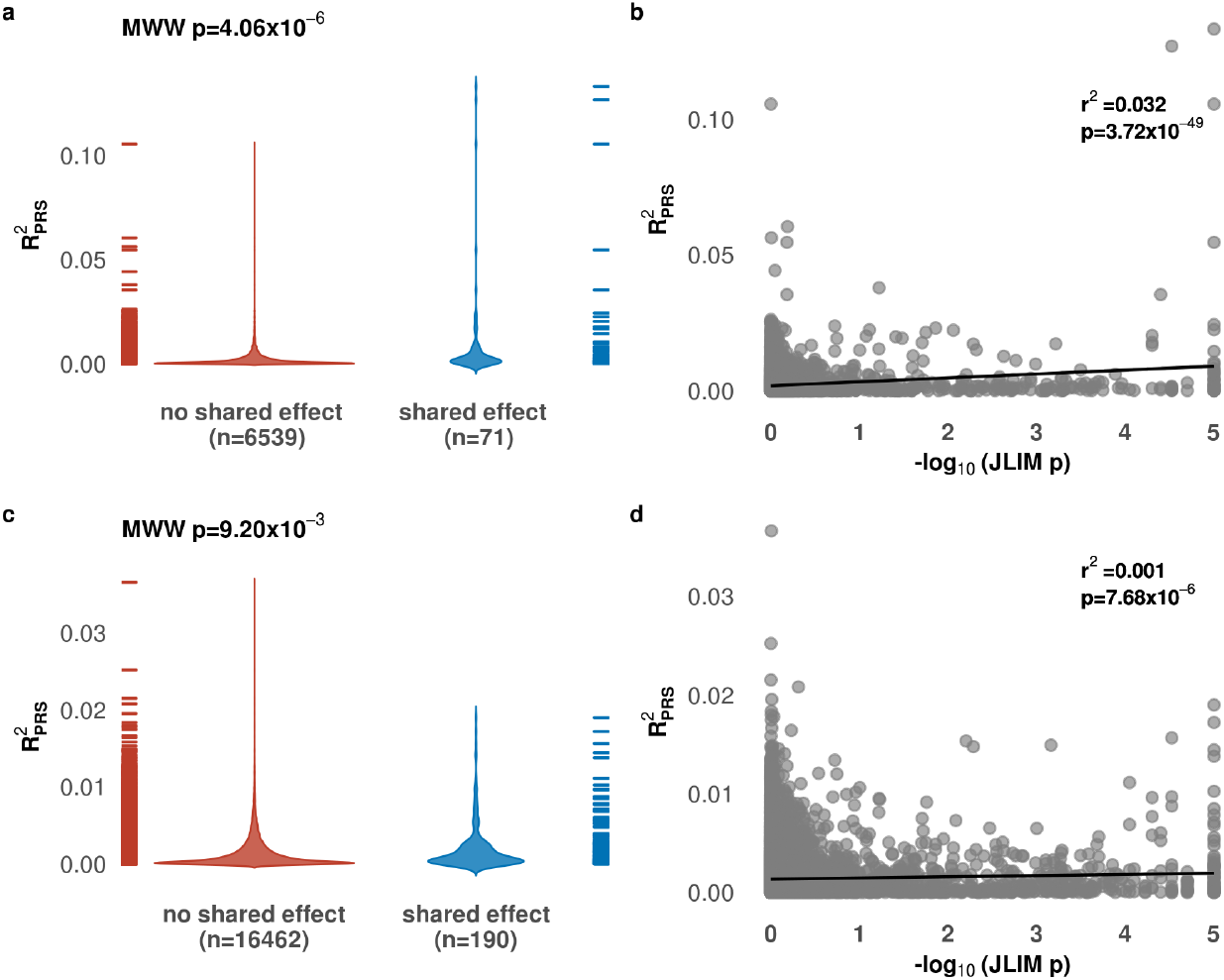
The immune phenotype variance explained by gene expression polygenic risk scores (PRS) is higher for trait pairs sharing associations. We considered 16,652 gene expression/immune trait pairs in our analysis. For each, we identified all independent variants meeting a threshold of association in the gene expression trait. We used these variants to calculate PRS for each individual in the Milieu Intérieur Project immune phenotype collection, and calculated the proportion of immune phenotype variance explained by these PRS 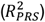. We found that gene expression PRS could explain significantly more immune trait variance for gene expression/immune trait pairs with a shared underlying genetic effect than for those that did not share an association. We saw this effect at different thresholds for selecting PRS instruments: (a) expression quantitative trait locus (eQTL) p < 5×10^−8^; and (c) eQTL p < 5×10^−5^. We also found that the proportion of variance explained was correlated to the overall strength of evidence for a shared (JLIM *p*-value) at these two selection thresholds (b, d). MWW = Mann-Whitney-Wilcoxon test.

**Table 1:** Expression quantitative trait locus (eQTL) variants explain more immune trait variance when they share an association. We considered 22,379 eQTL/immune trait pairs in our analysis. For each, we identified all independent variants meeting a threshold of association in the eQTL trait. We used these variants to calculate polygenic risk scores (PRS) for each individual in the Milieu Intérieur Project immune phenotype collection, and calculated the proportion of paired immune trait variance explained by that PRS 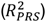 We compared 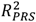 values for trait pairs with and without evidence for a shared underlying genetic effect using a Mann-Whithney-Wilcoxon test and determined the associations of JLIM *p*-values with 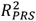 values and PRS *p*-values using univariate linear regression. To account for shared effects between some trait pairs but not others, we conditioned eQTL traits on the variant with the strongest JLIM *p*-value, and used the now conditionally independent eQTL data for the PRS calculation. Statistically significant results are shown in bold font.

| eQTL significance threshold ($p$-value) | Number of variants in gene expression trait PRS (mean, SD) | Number of trait pairs with shared genetic effect / Number of colocated trait pairs | Mean trait variance explained by gene expression PRS in trait pairs with shared vs. colocated associations | Difference in trait variance explained by gene expression PRS in shared vs. colocated associations (Mann-Whitney test $p$-value) | Evidence of shared association predicting immune trait variance explained by gene expression PRS (univariate regression $R^2$, $p$-value) |
| --- | --- | --- | --- | --- | --- |
| $5 \times 10^{-8}$ | 1.37, 0.69 | 71/6,539 | $1.00 \times 10^{-02} / 1.95 \times 10^{-03}$ | **$4.06 \times 10^{-06}$** | **0.032, $3.72 \times 10^{-49}$** |
| $1 \times 10^{-7}$ | 1.43, 0.75 | 77/7,177 | $9.42 \times 10^{-03} / 1.94 \times 10^{-03}$ | **$1.11 \times 10^{-05}$** | **0.025, $1.44 \times 10^{-41}$** |
| $5 \times 10^{-7}$ | 1.69, 0.97 | 118/10,128 | $6.03 \times 10^{-03} / 1.77 \times 10^{-03}$ | $5.25 \times 10^{-02}$ | **0.014, $2.73 \times 10^{-32}$** |
| $1 \times 10^{-6}$ | 2.00, 1.17 | 132/12,321 | $4.77 \times 10^{-03} / 1.69 \times 10^{-03}$ | $6.37 \times 10^{-02}$ | **0.009, $8.86 \times 10^{-26}$** |
| $5 \times 10^{-6}$ | 4.80, 2.28 | 189/16,293 | $2.76 \times 10^{-03} / 1.52 \times 10^{-03}$ | $7.06 \times 10^{-01}$ | **0.004, $4.19 \times 10^{-15}$** |
| $1 \times 10^{-5}$ | 8.54, 3.07 | 190/16,458 | $2.12 \times 10^{-03} / 1.45 \times 10^{-03}$ | $7.49 \times 10^{-01}$ | **0.002, $3.92 \times 10^{-08}$** |
| $5 \times 10^{-5}$ | 36.47, 6.29 | 190/16,462 | $2.00 \times 10^{-03} / 1.37 \times 10^{-03}$ | **$9.20 \times 10^{-03}$** | **0.001, $7.68 \times 10^{-06}$** |
| $1 \times 10^{-4}$ | 69.08, 8.75 | 190/16,462 | $1.90 \times 10^{-03} / 1.34 \times 10^{-03}$ | **$2.52 \times 10^{-03}$** | **0.001, $5.07 \times 10^{-04}$** |
| $1 \times 10^{-3}$ | 569.59, 26.20 | 190/16,462 | $1.70 \times 10^{-03} / 1.32 \times 10^{-03}$ | $3.03 \times 10^{-01}$ | **0.000, $4.55 \times 10^{-02}$** |
| $1 \times 10^{-2}$ | 4291.77, 72.84 | 190/16,462 | $1.50 \times 10^{-03} / 1.30 \times 10^{-03}$ | **$4.33 \times 10^{-02}$** | 0.000, $1.82 \times 10^{-01}$ |

To ensure our results were not due to selection artefacts induced by *p*-value thresholds, we also examined the correlation between all JLIM *p*-values and variance of immune phenotypes explained by the gene expression PRS, and found significant correlation (Table 1, Figure 3). Together, the results of the PRS analyses provide evidence for a stronger genetic correlation between colocalized gene expression and immune phenotypes if they share the same underlying genetic effect.

### Trait pairs sharing associations are more likely to be causally related

We next assessed evidence for causality directly with Mendelian randomization MR, again comparing trait pairs with a shared association to trait pairs with no evidence of shared associations. As our gene expression and immune phenotypes come from different cohorts, we used TSMR^28,29^. Statistical power in MR analyses is increased by selecting multiple variants as instruments^38^. This presents a problem when considering eQTLs as intermediate traits, because *cis*-acting effects often explain a large portion of phenotypic variance, and there is little power to detect *trans*-acting effects genome-wide^39–41^. To account for this power issue, we selected 16,631/16,652 trait pairs from above (190 with, and 16,441 without a shared association to an immune phenotype) where we found evidence for at least 2 independent variants associated with the gene expression trait (conditional *p* < 1×10^−5^). Using these variants as TSMR instruments, we found that the estimated causal effects of gene expression traits on immune phenotypes were higher in the 190 pairs with a shared association compared to the 16,441 non-sharing pairs (Figure 4a), but this difference was not significant. However, we observed overall correlation between JLIM *p*-values and the magnitude of the causal effect of gene expression traits on immune phenotypes (Figure 4b). These results suggest that trait pairs sharing an association are more likely to be causally related, and that this likelihood increases with increasing evidence that a shared association exists.

**Figure 4:**
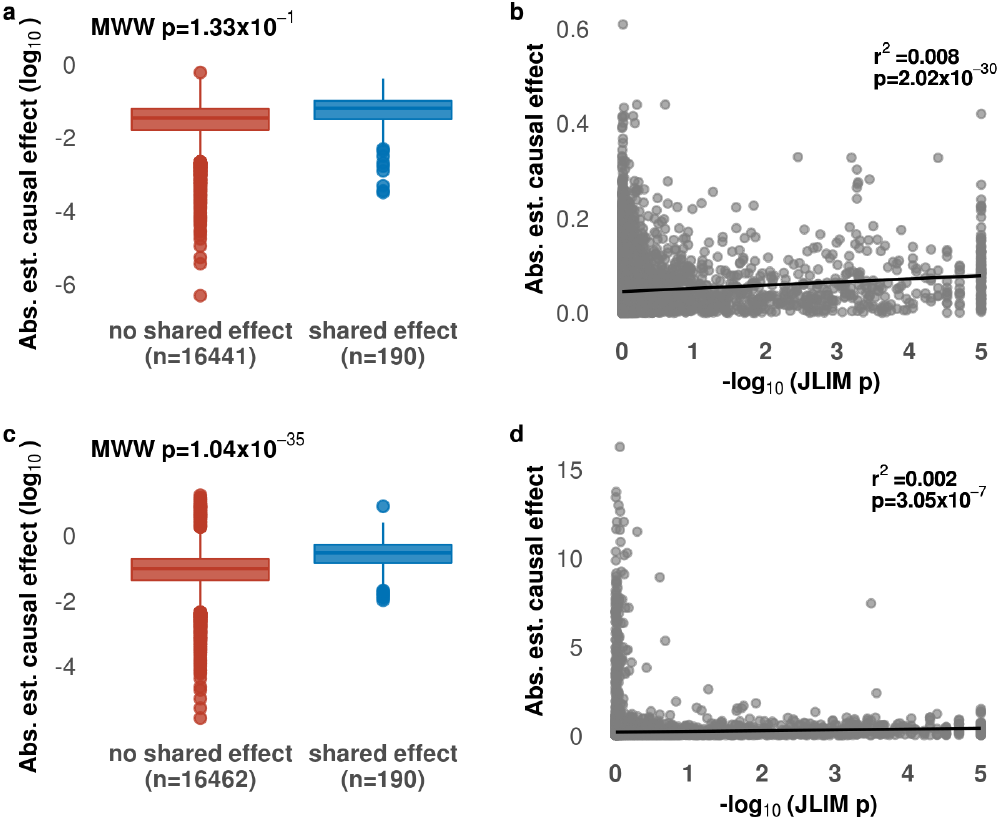
Trait pairs sharing an association are more likely to be causally related. We assessed evidence for causality between gene expression and immune cell traits using Mendelian randomization with cis-eQTL (expression quantitative trait loci) single nucleotide polymorpisms (SNPs) (a and b) and an expanded set of instruments including nearby genes (c and d). For the latter, we saw a significant level of increased evidence for causal effects in trait pairs with a shared effect (blue) compared to pairs without a shared effect (red) (c). We also saw an overall correlation between evidence for shared effects and the estimated causal effect size (b, d). MWW = Mann-Whitney-Wilcoxon test.

To address the limited number of instruments available for gene expression traits, we broadened our analysis to include other transcripts in the locus influenced by the same variants^30^. For each trait pair, we first identify variants independently associated with the gene expression trait, as above. We then ask if these variants are associated with any other transcript levels in the locus (within 1 megabase of the lead variant in the shared association test), and if so, identify all variants independently associated with those transcripts too. We thus gather a larger set of instruments for our MR analysis. We remove transcripts whose overall expression is highly correlated to the initial gene expression trait, to avoid over-estimating the effects of variants, as previously suggested^30^. We then perform the same inverse variance-weighted MR analysis as above, using the expanded instrument sets.

We found that gene expression traits explained a higher proportion of immune phenotype variance in the 190 pairs with shared associations, compared to the 15,462 other pairs (0.285 and 0.094, respectively; Figure 4c). As in our TSMR analysis, we saw a positive correlation between the strength of the JLIM *p*-value and the estimated causal effects of the gene expression traits on immune phenotypes (Figure 4d). Cumulatively, our results suggest that trait pairs with a shared association are more likely to be causally related than trait pairs that do not share an association.

## Discussion

In this work, we report that immune cell traits and gene expression traits share a small but significant set of associations, and that these point to interesting biological events with mechanistic implications. We then show that traits sharing a pleiotropic association tend to be causally related, rather than subject to horizontal pleiotropy. Thus, even though individual analyses may be under-powered, we are able to show in bulk that shared associations occur between causally related traits.

These shared associations can uncover previously unknown facets of immune cell biology. The shared effect between L-selectin expression and surface protein levels illustrates this principle: an allele that increases L-selectin transcript levels in neutrophils and monocytes also increases the mean fluorescence intensity of the L-selectin protein product CD62L expressed on neutrophil and eosinophil surface membranes. L-selectin is a cell adhesion molecule used by diverse immune cells to enter target organs by interacting with resident endothelial cells^42^. It is particularly important for the entry of naive T cells into secondary lymphoid tissues as part of the maturation process^43,44^. CD62L levels are thought to be predictive of treatment response in leukemia^45^ and risk of adverse events in multiple sclerosis therapy^46^; the effect of the L-selectin eQTL on protein levels could thus be misconstrued in a clinical setting.

We were struck by the number of shared effects between monocyte and neutrophil gene expression traits and T cell immune parameters. In these cases, the genes are either not detected at all, or show no evidence of an eQTL, in our T cell data (Figure 2). These results suggest that changes to gene expression in one cell type can have a direct effect on another population. Widespread cross-talk between immune cell subsets is a well-attested phenomenon, but to our knowledge this is the first time genetic mapping uncovers mechanisms of cellular coordination across cell subsets.

A major challenge in human genetics is translating genotype-phenotype associations into testable hypotheses of the underlying molecular, cellular and physiological events. Directly predicting the effect of a trait-associated variant remains challenging, especially for non-coding polymorphisms. This is further hampered by the limited resolution of fine mapping, so that in most cases we can only narrow an association signal to a group of variants over a genomic interval, all of which must be investigated, rather than pinpoint the exact causal variant. As variant function prediction ability is limited, direct experimentation is necessary, gradually uncovering molecular, cellular and ultimately physiological effects. Without prior information, a variety of outcomes across different cell types and conditions need to be assessed at each stage to uncover trait-relevant events. This approach is not scalable, so most translation efforts are necessarily conducted piecemeal.

One alternative approach to this bottleneck is to exploit pleiotropy across traits to generate molecular, cellular and physiological mechanism hypotheses, which can then be tested experimentally in a more focused way. This approach uses the fact that a variant associated with a physiological trait must act on molecular and cellular events; these are, by definition, also genetic traits as they are altered by a genetic variant, and the variant must have a pleiotropic effect on all these traits. We can thus compare, in unbiased fashion, many molecular traits (gene expression levels, in the present work) with many cellular traits (here, immunological parameters) to identify pleiotropic effects. Two broad approaches to this cross-trait pleiotropy have been articulated: the first is to look for shared heritability between two traits genome-wide^47,48^; and the second to look for shared effects at specific loci where we see genotype-phenotype associations, as we do in the present work. The latter is particularly suited to traits such as gene expression, where one variant often explains a large proportion of trait variance.

After identifying pairs of traits that share genetic effects (either locus-specific or genome-wide), it is tempting to immediately conclude that one trait directly causes the other. This conclusion is particularly appealing when considering traits across different levels of physiology, as we do here with gene expression and cellular measurements. Our results, however, suggest that simply observing association of different traits at the same genetic locus is not sufficient. Rather, we must look for direct evidence for causality between traits, and shared association mapping is a useful approach to do so. Ultimately, only with explicit proof of causality can we construct mechanistic hypotheses about trait physiology.

## Acknowledgements

This study makes use of data generated by the Blueprint Consortium. A full list of the investigators who contributed to the generation of the data is available from www.blueprint-epigenome.eu. Funding for the project was provided by the European Union’s Seventh Framework Programme (FP7/2007-2013) under grant agreement no 282510 – BLUEPRINT.

CG received a research fellowship from the Deutsche Forschungsgemeinschaft (DFG, German Research Foundation) for this project. She further received funding from the Hans und Klementia Langmatz-Stifung and the Hertie Network of Excellence in Clinical Neuroscience, not related to this study.

## Author contributions

All authors designed research. SC and SRS contributed software. CG and CC analyzed data and wrote the manuscript, which was approved by SC and SRS.

## References

1. He, X. et al. Sherlock: Detecting Gene-Disease Associations by Matching Patterns of Expression QTL and GWAS. Am. J. Hum. Genet. 92, 667–680 (2013).

2. Ongen, H. et al. Estimating the causal tissues for complex traits and diseases. Nat. Genet. 49, 1676–1683 (2017).

3. Akle, S. et al. Leveraging pleiotropy to discover and interpret GWAS results for sleep-associated traits. bioRxiv 832162 (2019) doi:10.1101/832162.

4. Voight, B. F. & Cotsapas, C. Human genetics offers an emerging picture of common pathways and mechanisms in autoimmunity. Curr. Opin. Immunol. 24, 552–557 (2012).

5. Wallace, C. et al. Statistical colocalization of monocyte gene expression and genetic risk variants for type 1 diabetes. Hum. Mol. Genet. 21, 2815–2824 (2012).

6. Wallace, C. Statistical Testing of Shared Genetic Control for Potentially Related Traits. Genet. Epidemiol. 37, 802–813 (2013).

7. Giambartolomei, C. et al. Bayesian Test for Colocalisation between Pairs of Genetic Association Studies Using Summary Statistics. PLOS Genet. 10, e1004383 (2014).

8. Zhu, Z. et al. Integration of summary data from GWAS and eQTL studies predicts complex trait gene targets. Nat. Genet. 48, 481–487 (2016).

9. Hormozdiari, F. et al. Colocalization of GWAS and eQTL Signals Detects Target Genes. Am. J. Hum. Genet. 99, 1245–1260 (2016).

10. Nica, A. C. et al. Candidate Causal Regulatory Effects by Integration of Expression QTLs with Complex Trait Genetic Associations. PLOS Genet. 6, e1000895 (2010).

11. Deng, Y. & Pan, W. A powerful and versatile colocalization test. PLOS Comput. Biol. 16, e1007778 (2020).

12. Chun, S. et al. Limited statistical evidence for shared genetic effects of eQTLs and autoimmune disease-associated loci in three major immune cell types. Nat. Genet. 49, 600–605 (2017).

13. Maurano, M. T. et al. Systematic Localization of Common Disease-Associated Variation in Regulatory DNA. Science 337, 1190–1195 (2012).

14. Trynka, G. et al. Disentangling the Effects of Colocalizing Genomic Annotations to Functionally Prioritize Non-coding Variants within Complex-Trait Loci. Am. J. Hum. Genet. 97, 139–152 (2015).

15. Gusev, A. et al. Partitioning heritability of regulatory and cell-type-specific variants across 11 common diseases. Am. J. Hum. Genet. 95, 535–552 (2014).

16. Patin, E. et al. Natural variation in the parameters of innate immune cells is preferentially driven by genetic factors. Nat. Immunol. 19, 302–314 (2018).

17. Chen, L. et al. Genetic Drivers of Epigenetic and Transcriptional Variation in Human Immune Cells. Cell 167, 1398–1414.e24 (2016).

18. R Core Team (2017). R: A Language and Environment for Statistical Computing. (R Foundation for Statistical Computing).

19. Dobin, A. et al. STAR: ultrafast universal RNA-seq aligner. Bioinforma. Oxf. Engl. 29, 15–21 (2013).

20. DeLuca, D. S. et al. RNA-SeQC: RNA-seq metrics for quality control and process optimization. Bioinformatics 28, 1530–1532 (2012).

21. Robinson, M. D. & Oshlack, A. A scaling normalization method for differential expression analysis of RNA-seq data. Genome Biol. 11, R25 (2010).

22. Durinck, S. et al. BioMart and Bioconductor: a powerful link between biological databases and microarray data analysis. Bioinforma. Oxf. Engl. 21, 3439–3440 (2005).

23. Durinck, S., Spellman, P. T., Birney, E. & Huber, W. Mapping identifiers for the integration of genomic datasets with the R/Bioconductor package biomaRt. Nat. Protoc. 4, 1184–1191 (2009).

24. Purcell, S. et al. PLINK: a tool set for whole-genome association and population-based linkage analyses. Am. J. Hum. Genet. 81, 559–575 (2007).

25. Stegle, O., Parts, L., Durbin, R. & Winn, J. A Bayesian framework to account for complex non-genetic factors in gene expression levels greatly increases power in eQTL studies. PLoS Comput. Biol. 6, e1000770 (2010).

26. Choi, S. W. & O’Reilly, P. F. PRSice-2: Polygenic Risk Score software for biobank-scale data. GigaScience 8, (2019).

27. International Schizophrenia Consortium et al. Common polygenic variation contributes to risk of schizophrenia and bipolar disorder. Nature 460, 748–752 (2009).

28. Pierce, B. L. & Burgess, S. Efficient design for Mendelian randomization studies: subsample and 2-sample instrumental variable estimators. Am. J. Epidemiol. 178, 1177–1184 (2013).

29. Lawlor, D. A. Commentary: Two-sample Mendelian randomization: opportunities and challenges. Int. J. Epidemiol. 45, 908–915 (2016).

30. Porcu, E. et al. Mendelian randomization integrating GWAS and eQTL data reveals genetic determinants of complex and clinical traits. Nat. Commun. 10, 3300 (2019).

31. Nicolae, D. L. et al. Trait-Associated SNPs Are More Likely to Be eQTLs: Annotation to Enhance Discovery from GWAS. PLoS Genet. 6, (2010).

32. Aguet, F. et al. Genetic effects on gene expression across human tissues. Nature 550, 204–213 (2017).

33. Aguet, F. et al. The GTEx Consortium atlas of genetic regulatory effects across human tissues. bioRxiv 787903 (2019) doi:10.1101/787903.

34. Smith, N. A., Coleman, C. B., Gewurz, B. E. & Rochford, R. CD21 (Complement Receptor 2) Is the Receptor for Epstein-Barr Virus Entry into T Cells. J. Virol. 94, e00428–20 (2020).

35. Smith, G. D. & Ebrahim, S. ‘Mendelian randomization’: can genetic epidemiology contribute to understanding environmental determinants of disease? Int. J. Epidemiol. 32, 1–22 (2003).

36. van Rheenen, W., Peyrot, W. J., Schork, A. J., Lee, S. H. & Wray, N. R. Genetic correlations of polygenic disease traits: from theory to practice. Nat. Rev. Genet. 20, 567–581 (2019).

37. Voight, B. F. et al. Plasma HDL cholesterol and risk of myocardial infarction: a mendelian randomisation study. The Lancet 380, 572–580 (2012).

38. Burgess, S., Butterworth, A. & Thompson, S. G. Mendelian randomization analysis with multiple genetic variants using summarized data. Genet. Epidemiol. 37, 658–665 (2013).

39. Brynedal, B. et al. Large-Scale trans-eQTLs Affect Hundreds of Transcripts and Mediate Patterns of Transcriptional Co-regulation. Am. J. Hum. Genet. 100, 581–591 (2017).

40. Stranger, B. E. & Raj, T. Genetics of human gene expression. Curr. Opin. Genet. Dev. 23, 627–634 (2013).

41. Grundberg, E. et al. Mapping cis-and trans-regulatory effects across multiple tissues in twins. Nat. Genet. 44, 1084–1089 (2012).

42. Ivetic, A., Hoskins Green, H. L. & Hart, S. J. L-selectin: A Major Regulator of Leukocyte Adhesion, Migration and Signaling. Front. Immunol. 10, 1068 (2019).

43. Pizcueta, P. & Luscinskas, F. W. Monoclonal antibody blockade of L-selectin inhibits mononuclear leukocyte recruitment to inflammatory sites in vivo. Am. J. Pathol. 145, 461–469 (1994).

44. Hogg, N. & Berlin, C. Structure and function of adhesion receptors in leukocyte trafficking. Immunol. Today 16, 327–330 (1995).

45. Sopper, S. et al. Reduced CD62L Expression on T Cells and Increased Soluble CD62L Levels Predict Molecular Response to Tyrosine Kinase Inhibitor Therapy in Early Chronic-Phase Chronic Myelogenous Leukemia. J. Clin. Oncol. Off. J. Am. Soc. Clin. Oncol. 35, 175–184 (2017).

46. Voortman, M. M. et al. The effect of disease modifying therapies on CD62L expression in multiple sclerosis. Mult. Scler. J. -Exp. Transl. Clin. 4, 2055217318800810 (2018).

47. Bulik-Sullivan, B. et al. An atlas of genetic correlations across human diseases and traits. Nat. Genet. 47, 1236–1241 (2015).

48. Brainstorm Consortium et al. Analysis of shared heritability in common disorders of the brain. Science 360, (2018).

