## Supplementary figures and tables for "Shared associations identify causal relationships between gene expression and immune cell phenotypes"

### Supplementary material

Figure S1: Analysis overview for identifying shared effects between gene expression and immunological traits 1

Figure S2: Six immune phenotypes were not normally distributed after rank transformation 2

Figure S3: Converting two quantitative immune phenotypes to dichotomous traits 3

Figure S4: Schematic of the analysis windows used to identify shared genetic effects between immune phenotypes and gene expression traits 4

Figure S5: Number of independent association signals per gene expression trait and locus 5

Figure S6: The power to detect shared associations depends on the strength of association to the secondary trait 6

Figure S7: Shared genetic association on chromosome 1 for L-selectin (CD62L) in eosinophils and SELL expression in neutrophils, monocytes and T cells. 7

Figure S8: Shared genetic association on chromosome 1 for CR2 expression in T cells and CD21 in three different B cell subpopulations. 8

Figure S9: Phenotypic variance of the expression of L-selectin in eosinophils explained by the polygenic risk scores (PRS) of four different genes with a shared genetic effect and by the combination of the PRS of these genes. 9

Table S1: Flow cytometry measurements for 166 immune phenotypes provided by the Milieu intérieur project 10

Table S2: Polygenic risk score (PRS) results without conditioning on shared genetic effects 16

### Figure S1: Analysis overview for identifying shared effects between gene expression and immunological traits

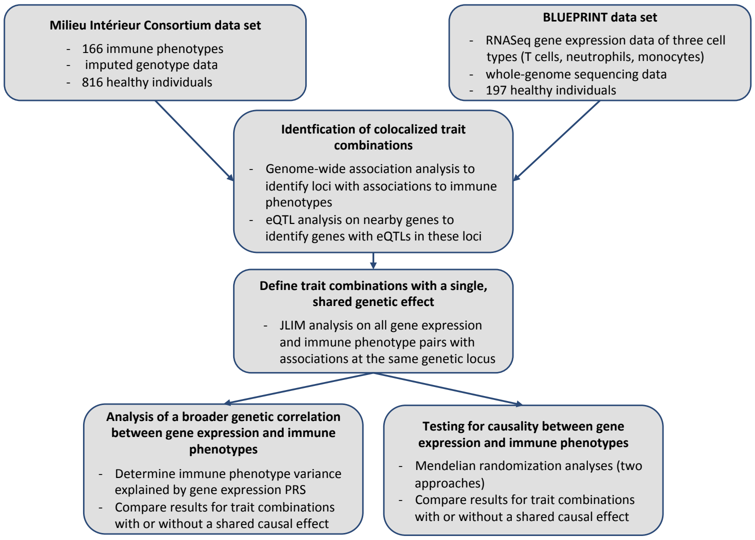


Abbreviations: eQTL = expression quantitative trait locus, PRS = polygenic risk score

**Figure S2: Six immune phenotypes were not normally distributed after rank transformation**

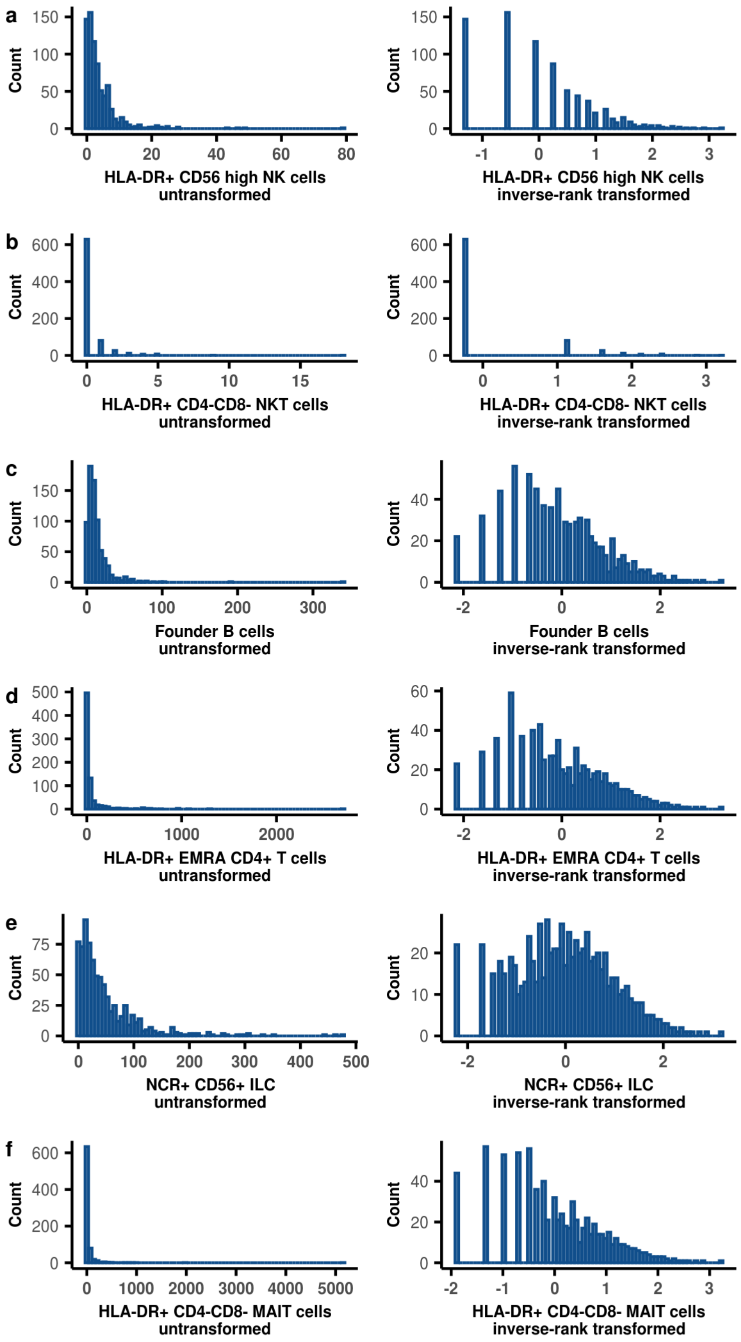


Six phenotypes were not normally distributed after rank transformation. After visual inspection, we removed two phenotypes - the number of HLA-DR+ CD56$^{high}$ NK cells (a) and the number of HLA-DR+ CD4-CD8- NKT cells (b) from further analysis and decided to binarize two other phenotypes - the number of founder B cells (c) and the number of HLA-DR positive CD4 positive EMRA T cells (e). We did not perform any other normalization for the two remaining phenotypes - the number of NCR+ CD56+ innate lymphoid cells (e) and the number of HLA-DR+ CD4-CD8- MAIT cells (f).

**Figure S3: Converting two quantitative immune phenotypes to dichotomous traits**

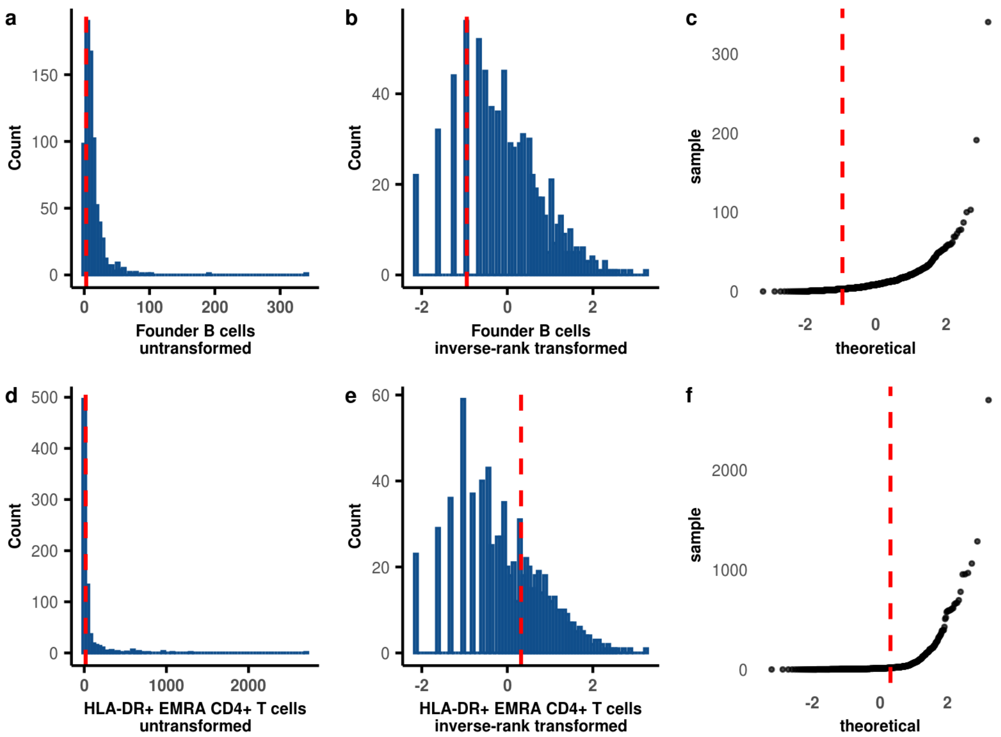


Two phenotypes - the number of founder B cells (panels a, b, and c) and the number of HLA-DR positive CD4 positive EMRA T cells (panels d, e, and f) - were binarized. Plots show the distribution of untransformed (a, d) as well as rank transformed (b, e) phenotypes and the quantile-quantile-plots (c, f). The thresholds for dichotomization (red dashed lines) were chosen as the point with the highest slope of the slope of the quantile-quantile plots.

**Figure S4: Schematic of the analysis windows used to identify shared genetic effects between immune phenotypes and gene expression traits**

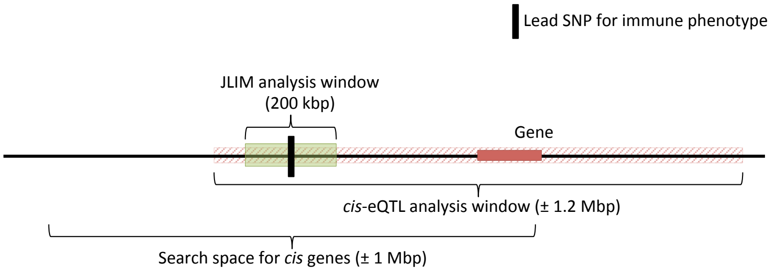


To identify effects shared between immune and gene expression phenotypes we first identified single nucleotide polymorphisms (SNPs) independently associated with an immune phenotype with association $p < 1\times10^{-5}$ (immune phenotype lead SNP). We then identified all genes with a transcription start site (TSS) within one megabase from that lead SNP. For each of these genes we performed a *cis*-expression quantiative trait locus (*cis*-eQTL) analysis to identify all indepently associated SNPs ($p < 1\times10^{-3}$) within  1.2 megabase from the TSS. For each gene with an eQTL with $p < 1\times10^{-3}$ within 200 kilobase from the immune phenotype lead SNP (JLIM analysis window) we then performed JLIM analysis on all SNPs within the JLIM analysis window to test for evidence of a shared genetic effect between the immune phenotype and gene expression traits.

**Figure S5: Number of independent association signals per gene expression trait and locus**

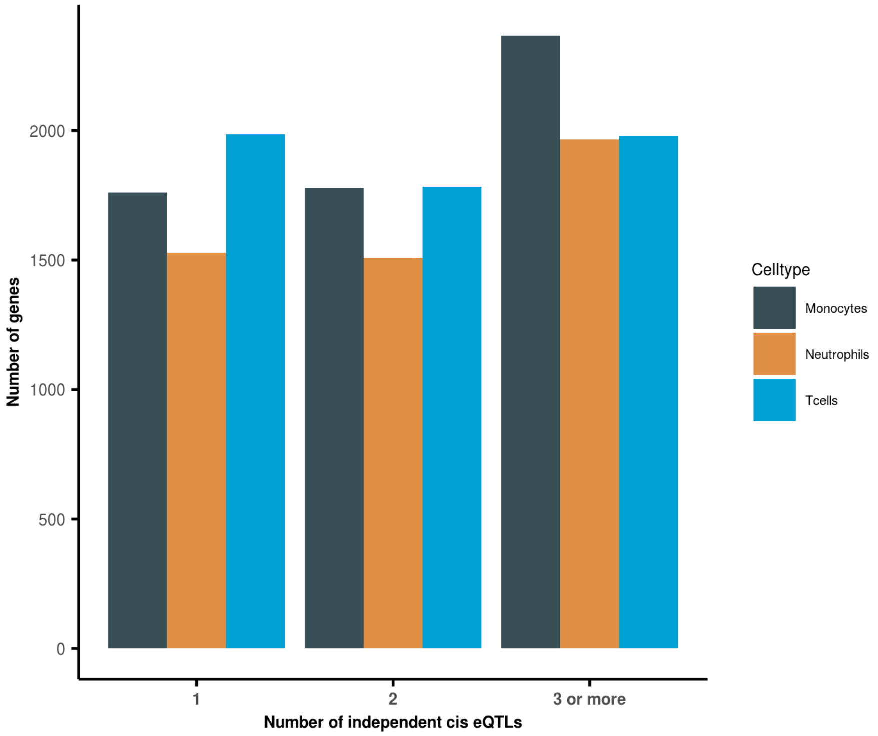


We performed *cis*-expression quantiative trait locus (*cis*-eQTL) analysis on 7,365 genes with a transcription start site (TSS) within 1 megabase from an immune phenotype lead SNP. For 5,906, 5,004 and 5,748 genes we found at least one *cis*-eQTL (within 1.2 megabases from the TSS) in monocytes (gray), neutrophils (yellow) and T cells (blue), respectively. For about two-thirds of these genes we could identify more than one independent *cis*-eQTL.

**Figure S6: The power to detect shared associations depends on the strength of association to the secondary trait**

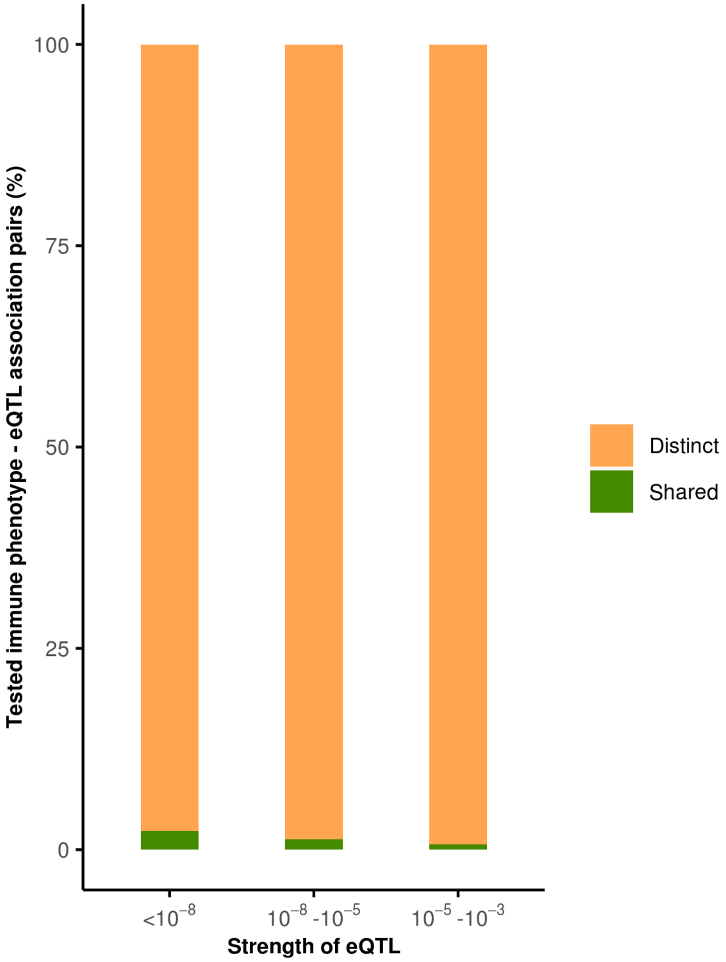


We find evidence that in most cases of colocalized expression quantitative trait loci (eQTLs) and immune phenotypes associations the genetic effects underlying these associations are distinct (orange). The proportion of shared genetic effects (green) we are able to detect depends on the strength of the eQTL association and is highest for strong eQTLs with association $p < 10^{-8}$.

**Figure S7: Shared genetic association on chromosome 1 for L-selectin (CD62L) in eosinophils and SELL expression in neutrophils, monocytes and T cells.**

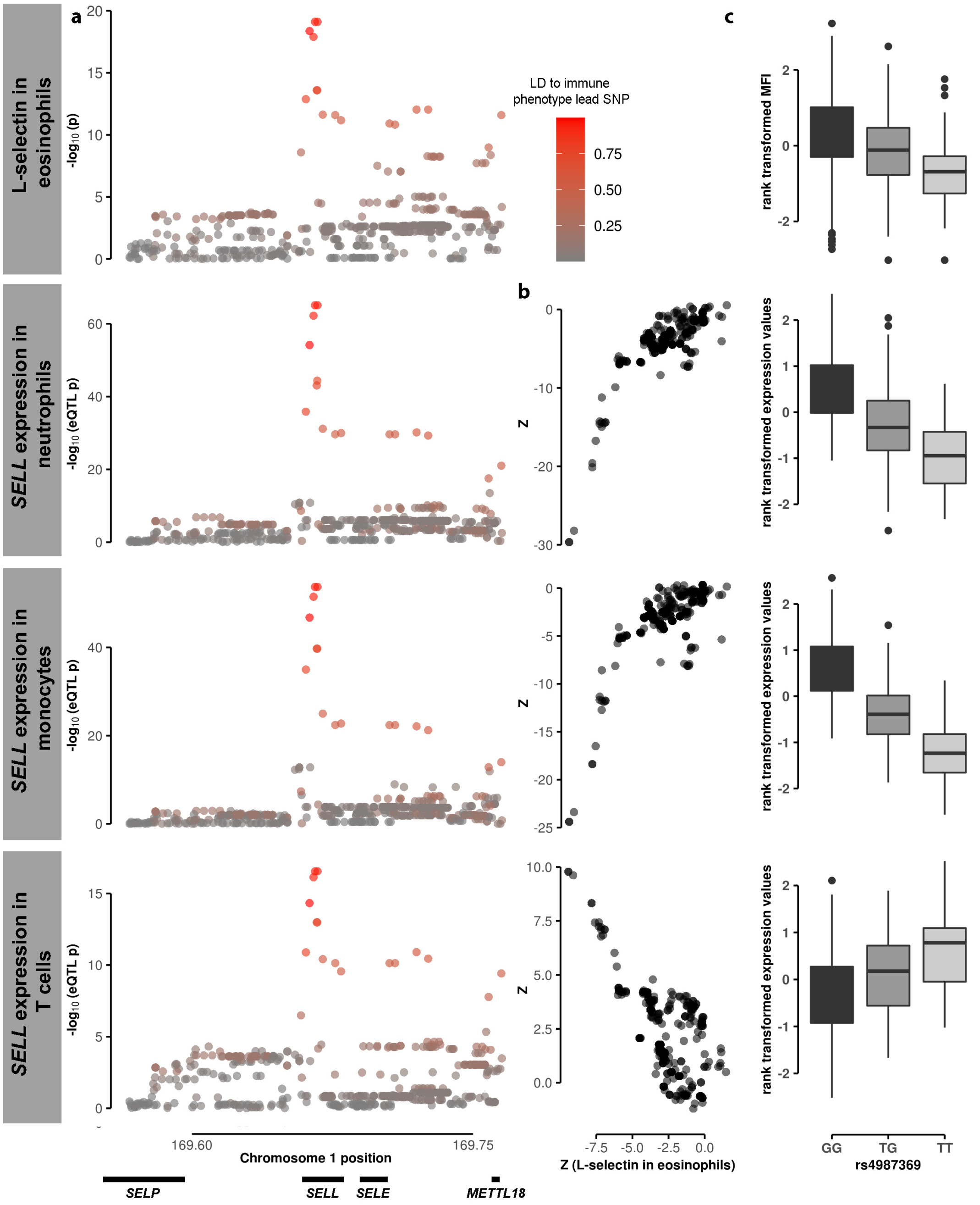


We find a shared genetic association with mean fluorescence intensity (MFI) of L-selectin (CD62L) in neutrophils and with *SELL* expression in neutrophils, monocytes and T cells (a). The association Z statistics of the MFI of L-selectin in neutrophils and *SELL* expression in the three different cell types are strongly correlated (b). (c) shows the MFI of L-selectin and *SELL* expression in the three different cell types (both rank transformed) per genotype of the lead variant rs4987369.

**Figure S8: Shared genetic association on chromosome 1 for CR2 expression in T cells and CD21 in three different B cell subpopulations.**

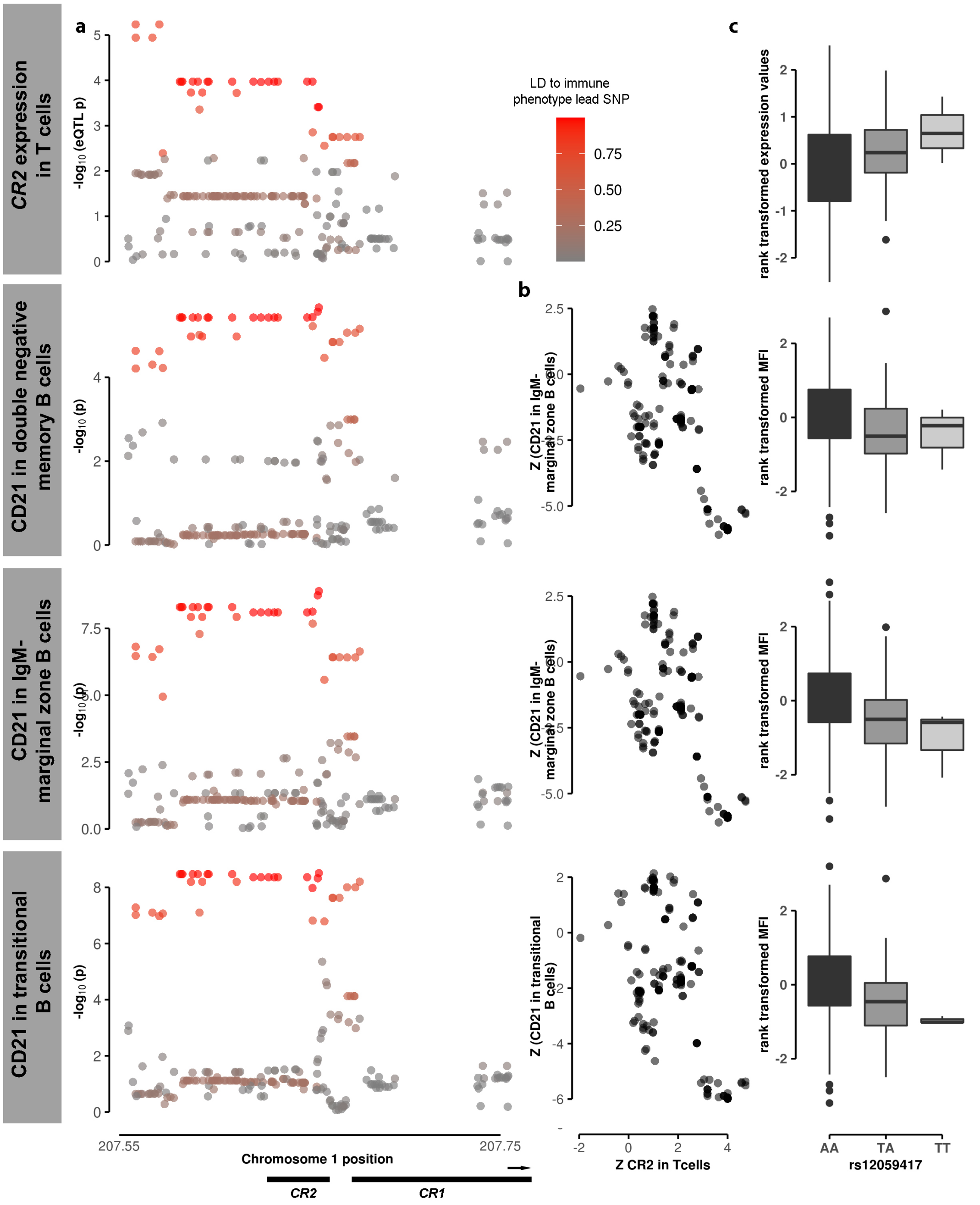


The association signal of the mean fluorescence intensity (MFI) of CD21 in three different B cell subpopulations - double negative memory B cells, IgM- marginal zone B cells and transitional B cells - at a genetic locus on chromosome 1 is consistent with the association signal to *CR2* expression in T cells (a). The association Z statistics of the MFI of CD21 in the three B cell subpopulations and *CR2* expression in T cells are strongly correlated (b). (c) shows the MFI of CD21 in the three B cell subpopulation and the level of *CR2* expression in T cells (all rank transformed) per genotype of the lead variant rs12059417.

**Figure S9: Phenotypic variance of the expression of L-selectin in eosinophils explained by the polygenic risk scores (PRS) of four different genes with a shared genetic effect and by the combination of the PRS of these genes.**

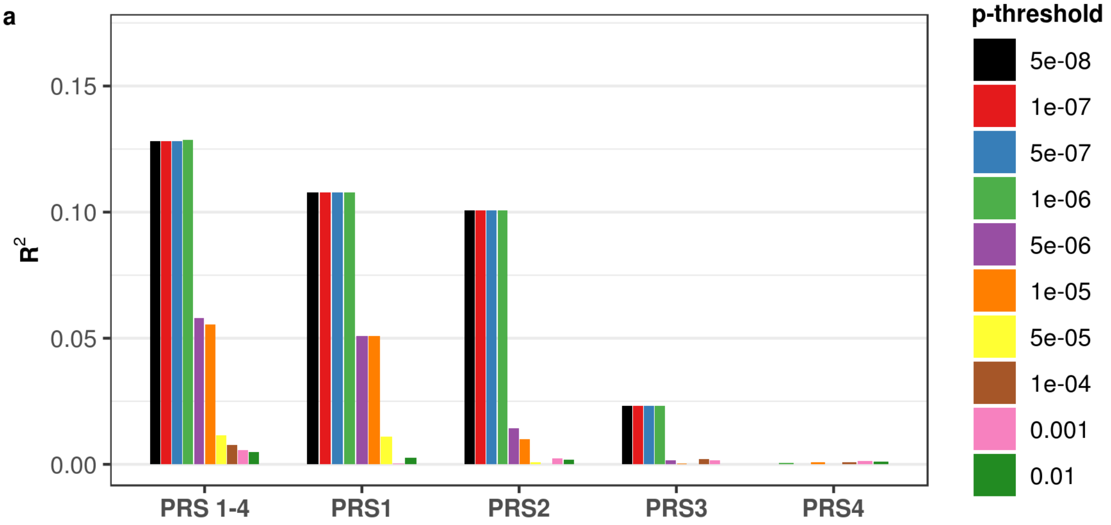


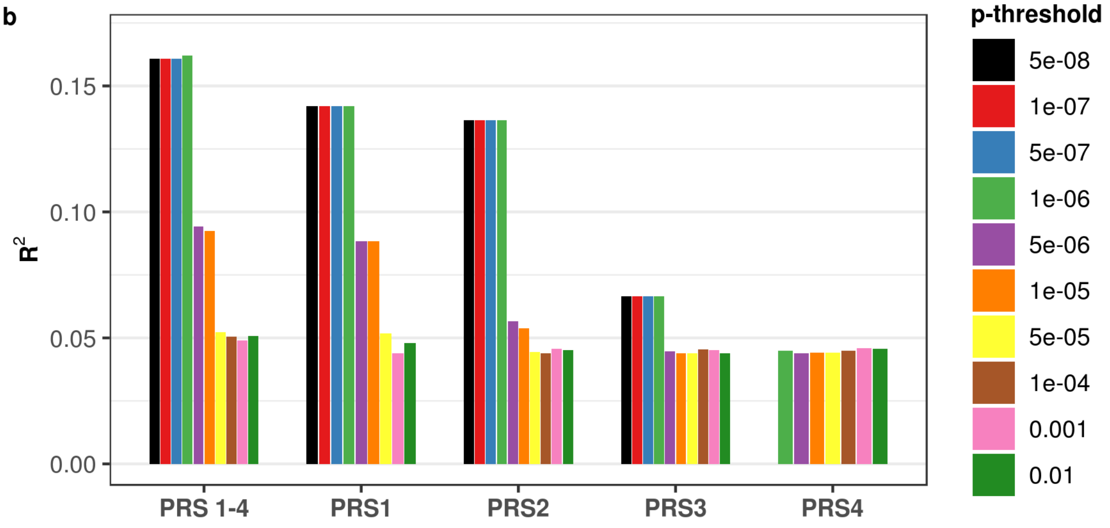


The phenotypic variance ($R^2$) of the expression of L-selectin (CD62L) in eosinophils explained by single polygenic risk scores (PRS) for four different genes with evidence for a shared genetic effect with the immune phenotype - SELL (PRS1), SELP (PRS2), TNF786 (PRS3) and the pseudogene EEF1A1P2 (PRS4) as well as by the combination of all four PRS (PRS 1-4) has been calculated by linear regression. Regression models were run without any other predictors (A) or including age, sex, CMV status and smoking status as well as the first five principal components in the regresion models (B). The phenotypic variance explained was highest when including the PRS scores of all four genes with evidence for a shared genetic effect with the expression of L-selectin (CD62L) in eosinophils into the regression models. The selection of single nucleotide polymorphisms (SNPs) included in the PRS calculation was based on ten eQTL p-thresholds (see color legend).

**Table S1: Flow cytometry measurements for 166 immune phenotypes provided by the Milieu intérieur project**

| Immunophenotype | Measure type |
|---|---|
| CD16hi NK cells | Absolute count |
| CD56hi NK cells | Absolute count |
| CD69+ CD16hi NK cells | Absolute count |
| CD69+ CD56hi NK cells | Absolute count |
| CD8a+ CD16hi NK cells | Absolute count |
| CD8a+ CD56hi NK cells | Absolute count |
| HLA-DR+ CD56hi NK cells | Absolute count |
| NK cells | Absolute count |
| CD16 in CD16hi NK cells | MFI |
| CD16 in CD56hi NK cells | MFI |
| CD69 in CD16hi NK cells | MFI |
| CD69 in CD56hi NK cells | MFI |
| CD69 in CD69+ CD16+ NK cells | MFI |
| CD69 in CD69+ CD56+ NK cells | MFI |
| CD69 in CD8a+ CD16+ NK cells | MFI |
| CD69 in CD8a+ CD56+ NK cells | MFI |
| CD69 in HLA-DR+ CD16+ NK cells | MFI |
| CD8a in CD16hi NK cells | MFI |
| CD8a in CD56hi NK cells | MFI |
| CD8a in CD69+ CD16+ NK cells | MFI |
| CD8a in CD69+ CD56+ NK cells | MFI |
| CD8a in CD8a+ CD16+ NK cells | MFI |
| CD8a in CD8a+ CD56+ NK cells | MFI |
| CD8a in HLA-DR+ CD16+ NK cells | MFI |
| HLA-DR in CD16hi NK cells | MFI |

| | |
|---|---|
| HLA-DR in CD56hi NK cells | MFI |
| HLA-DR in CD69+ CD16+ NK cells | MFI |
| HLA-DR in CD69+ CD56+ NK cells | MFI |
| HLA-DR in CD8a+ CD16+ NK cells | MFI |
| HLA-DR in CD8a+ CD56+ NK cells | MFI |
| HLA-DR in HLA-DR+ CD16+ NK cells | MFI |
| HLA-DR in HLA-DR+ CD56+ NK cells | MFI |
| NKp46 in NK cells | MFI |
| ratio of CD16 MFI in CD16hi to CD56hi NK cells | Ratio |
| ratio of CD16hi to CD56hi NK cells | Ratio |
| CD14hi monocytes | Absolute count |
| CD16hi monocytes | Absolute count |
| monocytes | Absolute count |
| CD16 in CD14hi monocytes | MFI |
| CD16 in CD16hi monocytes | MFI |
| basophils | Absolute count |
| eosinophils | Absolute count |
| neutrophils | Absolute count |
| CD16 in basophils | MFI |
| CD16 in eosinophils | MFI |
| CD16 in neutrophils | MFI |
| CD203c in basophils | MFI |
| CD32 in basophils | MFI |
| CD62L in eosinophils | MFI |
| CD62L in neutrophils | MFI |
| FcεRI in basophils | MFI |
| FcεRI in eosinophils | MFI |
| FcεRI in neutrophils | MFI |
| cDC1 | Absolute count |
| cDC3 | Absolute count |

| | |
|---|---|
| pDC | Absolute count |
| CD86 in CD14hi monocytes | MFI |
| CD86 in cDC1 | MFI |
| CD86 in cDC3 | MFI |
| CD86 in pDC | MFI |
| HLA-DR in CD14hi monocytes | MFI |
| HLA-DR in cDC1 | MFI |
| HLA-DR in cDC3 | MFI |
| HLA-DR in pDC | MFI |
| CD161+ ILC3 | Absolute count |
| CD56+ ILC | Absolute count |
| ILC | Absolute count |
| ILC1 | Absolute count |
| ILC2 | Absolute count |
| ILC3 | Absolute count |
| NCR- CD56+ ILC | Absolute count |
| NCR+ CD56+ ILC | Absolute count |
| CD127 in ILC1 | MFI |
| CD161 in ILC1 | MFI |
| CD161 in ILC3 | MFI |
| CD4+ T cells | Absolute count |
| CD8b- CD4- T cells | Absolute count |
| CD8b+ T cells | Absolute count |
| CM CD4+ T cells | Absolute count |
| CM CD8+ T cells | Absolute count |
| EM CD4+ T cells | Absolute count |
| EM CD8+ T cells | Absolute count |
| EMRA CD4+ T cells | Absolute count |
| EMRA CD8+ T cells | Absolute count |
| HLA-DR+ CM CD4+ T cells | Absolute count |

| | |
|---|---|
| HLA-DR+ CM CD8+ T cells | Absolute count |
| HLA-DR+ EM CD8+ T cells | Absolute count |
| HLA-DR+ EMRA CD4+ T cells | Absolute count |
| HLA-DR+ EMRA CD8+ T cells | Absolute count |
| HLA-DR+ in EM CD4+ T cells | Absolute count |
| naïve CD4+ T cells | Absolute count |
| naïve CD8+ T cells | Absolute count |
| T cells | Absolute count |
| CCR7 in CM CD4+ T cells | MFI |
| CCR7 in CM CD8+ T cells | MFI |
| CCR7 in EM CD4+ T cells | MFI |
| CCR7 in EM CD8+ T cells | MFI |
| CCR7 in EMRA CD4+ T cells | MFI |
| CCR7 in EMRA CD8+ T cells | MFI |
| CCR7 in naïve CD4+ T cells | MFI |
| CCR7 in naïve CD8+ T cells | MFI |
| CD4:CD8 ratio | Ratio |
| activated Treg | Absolute count |
| conventional T cells | Absolute count |
| memory Treg | Absolute count |
| naïve Treg | Absolute count |
| Treg cells | Absolute count |
| ICOS in activated Treg | MFI |
| ICOS in memory Treg | MFI |
| ICOS in naïve Treg | MFI |
| CD4- CD8- MAIT cells | Absolute count |
| CD4- CD8- NKT cells | Absolute count |
| CD8+ MAIT cells | Absolute count |
| HLA-DR+ CD4- CD8- MAIT cells | Absolute count |
| HLA-DR+ CD4- CD8- NKT cells | Absolute count |

| | |
|---|---|
| MAIT cells | Absolute count |
| NKT in T cells | Absolute count |
| γδ+ T cells | Absolute count |
| B cells | Absolute count |
| CD21- CD27- B cells | Absolute count |
| CD21- CD27+ B cells | Absolute count |
| CD24hi memory B cells | Absolute count |
| CD24low memory B cells | Absolute count |
| double negative memory B cells | Absolute count |
| founder B cells | Absolute count |
| germinal center B cells | Absolute count |
| IgM- marginal zone B cells | Absolute count |
| IgM+ marginal zone B cells | Absolute count |
| marginal zone B cells | Absolute count |
| memory B cells | Absolute count |
| naïve B cells | Absolute count |
| transitional B cells | Absolute count |
| CD19 in B cells | MFI |
| CD21 in B cells | MFI |
| CD21 in CD24hi memory B cells | MFI |
| CD21 in CD24int memory B cells | MFI |
| CD21 in CD24low memory B cells | MFI |
| CD21 in double negative memory B cells | MFI |
| CD21 in germinal center B cells | MFI |
| CD21 in IgM- marginal zone B cells | MFI |
| CD21 in IgM+ marginal zone B cells | MFI |
| CD21 in marginal zone B cells | MFI |
| CD21 in memory B cells | MFI |
| CD21 in naïve B cells | MFI |
| CD21 in transitional B cells | MFI |

| | |
|---|---|
| CD24 in B cells | MFI |
| CD24 in double negative memory B cells | MFI |
| CD24 in germinal center B cells | MFI |
| CD24 in IgM- marginal zone B cells | MFI |
| CD24 in IgM+ marginal zone B cells | MFI |
| CD24 in marginal zone B cells | MFI |
| CD24 in memory B cells | MFI |
| CD24 in naïve B Cells | MFI |
| CD27 in B cells | MFI |
| CD38 in B cells | MFI |
| CD38 in double negative memory B cells | MFI |
| CD38 in germinal center B cells | MFI |
| CD38 in memory B cells | MFI |
| IgD in B cells | MFI |
| IgM in B cells | MFI |
| CCR6+ CD4+ T cells | Absolute count |
| CCR6+ CD8+ T cells | Absolute count |
| CRTh2+ CD4+ T cells | Absolute count |
| CXCR5+ CD4+ T cells | Absolute count |
| CD45+ cells | Absolute count |
| total number of cells | Absolute count |

**Table S2: Polygenic risk score (PRS) results without conditioning on shared genetic effects**

| eQTL significance threshold (*p*-value) | Number of variants in gene expression trait PRS (mean, SD) | Number of trait pairs with shared genetic effect / Number of colocalized trait pairs | Mean trait variance explained by gene expression PRS in trait pairs with shared vs. colocalized associations | Difference in trait variance explained by gene expression PRS in shared vs. colocalized associations (Mann-Whitney test, *p*-value) | Evidence of shared association predicting immune trait variance explained by gene expression PRS (univariate regression $R^2$, *p*-value) |
|---|---|---|---|---|---|
| $5\times10^{-8}$ | 1.42, 0.75 | 118/8,134 | $2.52\times10^{-02}$ / $2.43\times10^{-03}$ | **$3.56\times10^{-35}$** | **0.160, $< 1\times10^{-100}$** |
| $1\times10^{-7}$ | 1.47, 0.81 | 125/8,781 | $2.54\times10^{-02}$ / $2.42\times10^{-03}$ | **$2.16\times10^{-36}$** | **0.157, $< 1\times10^{-100}$** |
| $5\times10^{-7}$ | 1.79, 1.05 | 155/11,380 | $1.95\times10^{-02}$ / $2.18\times10^{-03}$ | **$2.27\times10^{-30}$** | **0.120, $< 1\times10^{-100}$** |
| $1\times10^{-6}$ | 2.11, 1.26 | 163/13,237 | $1.76\times10^{-02}$ / $2.01\times10^{-03}$ | **$7.49\times10^{-32}$** | **0.120, $< 1\times10^{-100}$** |
| $5\times10^{-6}$ | 5.06, 2.36 | 190/16,306 | $1.06\times10^{-02}$ / $1.68\times10^{-03}$ | **$7.13\times10^{-24}$** | **0.070, $< 1\times10^{-100}$** |
| $1\times10^{-5}$ | 8.87, 3.15 | 190/16,460 | $7.82\times10^{-03}$ / $1.57\times10^{-03}$ | **$3.66\times10^{-24}$** | **0.053, $< 1\times10^{-100}$** |
| $5\times10^{-5}$ | 36.84, 6.45 | 190/16,462 | $3.73\times10^{-03}$ / $1.48\times10^{-03}$ | **$2.48\times10^{-08}$** | **0.009, $2.48\times10^{-35}$** |
| $1\times10^{-4}$ | 69.47, 8.86 | 190/16,462 | $2.82\times10^{-03}$ / $1.31\times10^{-03}$ | **$6.44\times10^{-06}$** | **0.004, $2.04\times10^{-16}$** |
| $1\times10^{-3}$ | 570.24, 26.17 | 190/16,462 | $1.72\times10^{-03}$ / $1.34\times10^{-03}$ | $5.26\times10^{-02}$ | 0.000, $1.36\times10^{-01}$ |
| $1\times10^{-2}$ | 4292.85, 73.14 | 190/16,462 | $1.41\times10^{-03}$ / $1.33\times10^{-03}$ | $8.38\times10^{-02}$ | 0.000, $3.22\times10^{-01}$ |

We considered 22,379 gene expression/immune trait pairs in our analysis. For each, we identified all independent variants meeting a threshold of association in the eQTL trait. We used these variants to calculate polygenic risk scores (PRS) for each individual in the Milieu Intérieur Project immune phenotype collection, and calculated the proportion of paired immune trait variance explained by that PRS ($R^2_{PRS}$). We compared $R^2_{PRS}$ values for trait pairs with and without evidence for a shared underlying genetic effect using a Mann-Whithney-Wilcoxon test and determined the associations of JLIM *p*-values with $R^2_{PRS}$ values and PRS *p*-values using univariate linear regression.
